# Structure Learning for Hierarchical Regulatory Networks

**DOI:** 10.1101/2021.05.27.446022

**Authors:** Anthony Federico, Joseph Kern, Xaralabos Varelas, Stefano Monti

## Abstract

Network analysis offers a powerful technique to model the relationships between genes within biological regulatory networks. Inference of biological network structures is often performed on high-dimensional data, yet is hindered by the limited sample size of high throughput “omics” data typically available. To overcome this challenge, we exploit known organizing principles of biological networks that are sparse, modular, and likely share a large portion of their underlying architecture. We present *SHINE* - **S**tructure Learning for **Hi**erarchical **Ne**tworks - a framework for defining data-driven structural constraints and incorporating a shared learning paradigm for efficiently learning multiple networks from high-dimensional data. We show through simulations *SHINE* improves performance when relatively few samples are available and multiple networks are desired, by reducing the complexity of the graphical search space and by taking advantage of shared structural information. We evaluated *SHINE* on TCGA Pan-Cancer data and found learned tumor-specific networks exhibit expected graph properties of real biological networks, recapture previously validated interactions, and recapitulate findings in literature. Application of *SHINE* to the analysis of subtype-specific breast cancer networks identified key genes and biological processes for tumor maintenance and survival as well as potential therapeutic targets for modulating known breast cancer disease genes.

## INTRODUCTION

Biological networks can model functional relationships at different cellular levels – genes, proteins, metabolites – and can be integrated to depict system-wide connectivity. Gene regulatory network (GRN) reconstruction aimed at inferring putative mechanistic interactions associated with disease phenotypes can support the identification of drivers of disease severity and treatment response. Importantly, changes in network connectivity across experimental conditions or phenotypes may help pinpoint important context-specific regulators or mediators, and inform functional experiments aimed at elucidating mechanisms of action (MOAs), targetable vulnerabilities, and resistance to treatment.

GRNs represent the underlying structure that dictate functional properties of biological systems. The dependencies between genes provide insight into the regulatory mechanisms that drive biological phenomena. Markov networks - undirected graphical models satisfying the Markov property, i.e., capable of distinguishing between direct and indirect dependencies – support a semantically rich representation of GRNs. Often studies forgo Markov graphical modeling due to the unavailability of the large number of samples required for their inference in high-dimensional domains. Comparing networks across multiple phenotypic groups or time points further dilutes already limited sample sizes, presenting an even greater challenge for network inference from omics data.

Gene connectivity is often inferred through pairwise correlation across genome-wide expression data. While a co-expression matrix is straightforward to compute, it fails to distinguish direct from indirect dependencies between genes. To infer direct effects, one must consider the partial correlation of genes, that is the correlation of two genes conditioned on all other genes. To that end, in linear settings one must compute the partial correlation or precision matrix Ω, which is intractable in high dimensional domains. Conventionally, the maximum-likelihood method is used to estimate Ω. However, when *n* < *p* – i.e., when the number of variables *p* is larger than the sample size *n*, as in the case of typical genome-wide omics data – the estimation of Ω becomes unstable and inaccurate.

Several regularization approaches have been developed to improve the estimation of Ω, such as the graphical lasso and its extensions^1,2^. An alternative method for learning Markov networks is a Bayesian approach to search for likely graphs that encode the data^3,4^. Bayesian structure learning also enables the incorporation of prior beliefs in the graphical search. However, all these solutions are still insufficient to achieve accurate inference when n≪p. In this report, we develop and evaluate a multi-pronged approach to Markov network reconstruction that combines Bayesian inference with constraint learning – to incorporate well-established modular properties of biological networks – and with shared learning – to pool information across networks representing related domains (e.g., tumor subtypes). In doing so, we limit the search complexity and increase the equivalent sample size for graphical modeling of high dimensional biological data.

Due to an exponentially expanding solution space in structure learning – with 2^*p*(*p*-1)/2^ possible graphs to choose from – finding appropriate constraints to limit the number of graphs to consider can drastically improve inference, especially for high dimensional models. More specifically, as entities in the cell – such as genes, proteins, metabolites – form co-regulated modules expected to share regulatory programs^5^, we use this module structure to form a high-level representation of an integrated regulatory space by estimating the modules’ interdependencies. We can thus reduce the complexity of the graphical search space by first identifying module-based constraints, and by then ruling out unlikely inter-module gene interactions. Additionally, when learning multiple networks representing specializations of a common domain, it is important to consider evidence that their topology is partially conserved – even between evolutionarily distinct species – particularly for genes involved in core biological processes^6^. Thus, a shared learning approach where data is shared between related phenotypes can be used to address underpowered analysis of high dimensional biological datasets with a small number of samples^7^.

## RESULTS

### Constrained Learning Limits the Structure Search Space

A well-accepted approach for reducing the complexity of gene-based models is to reduce the dimensionality of the data to co-expression modules. Given a *p x n* gene expression matrix *X*, by taking advantage of the zero-order correlations between expression profiles *x_i_*, and *x_j_* across *n* samples, genes can be organized into co-expression modules. This is based on the principle that complex biological systems have a modular structure and gene regulatory networks carry out biological functions through units of coordinated expression^8–10^. To this end, many methods have emerged to detect network modules based on gene co-expression^11^. Accounting for network modularity before Markov network inference allows one to first consider genes likely to be interacting based on co-expression similarity, which can be computed with few samples. The simplest approach is to detect mutually exclusive - or isolated - modules and to then constrain the graphical search to intra-modular interactions. A range of approaches can then be defined, based on varying criteria of stringency in constraining the inter-modular relationships to be modeled.

### Isolated Modules

Using methods popularized in Weighted Correlation Network Analysis (WGCNA)^12^ and following consistent notation^13^, genes are clustered by their co-expression similarity *S_ij_* – measured by the absolute value of the biweight midcorrelation^14^ coefficient: *s_i,j_* = |*bicor*(*x_i_, x_j_*)| as well as a soft thresholding value *β* which pushes spurious correlations to zero, resulting in a symmetric *p x p* weighted adjacency matrix 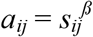. Co-expression modules are detected using hierarchical clustering of a topological overlap dissimilarity transformation *d_i,j_* of *a_ij_* resulting in *Q* modules. These modules represent interacting functional units that serve as a reasonable first order approximation of the larger biological network organization.

Here, we use them as a structural constraint when estimating the full graph (Fig. 1A), where the graphical search only considers the interaction of genes *i* and *j* if they belong to the same module (Fig. 1B). However, ignoring inter-module interactions imposes a strict constraint and yields a disconnected global structure, which is contrary to our understanding that biological processes are interconnected^15^. Therefore, we consider inter-modular interactions by finding genes coexpressed across two or more modules.

**Figure 1.**
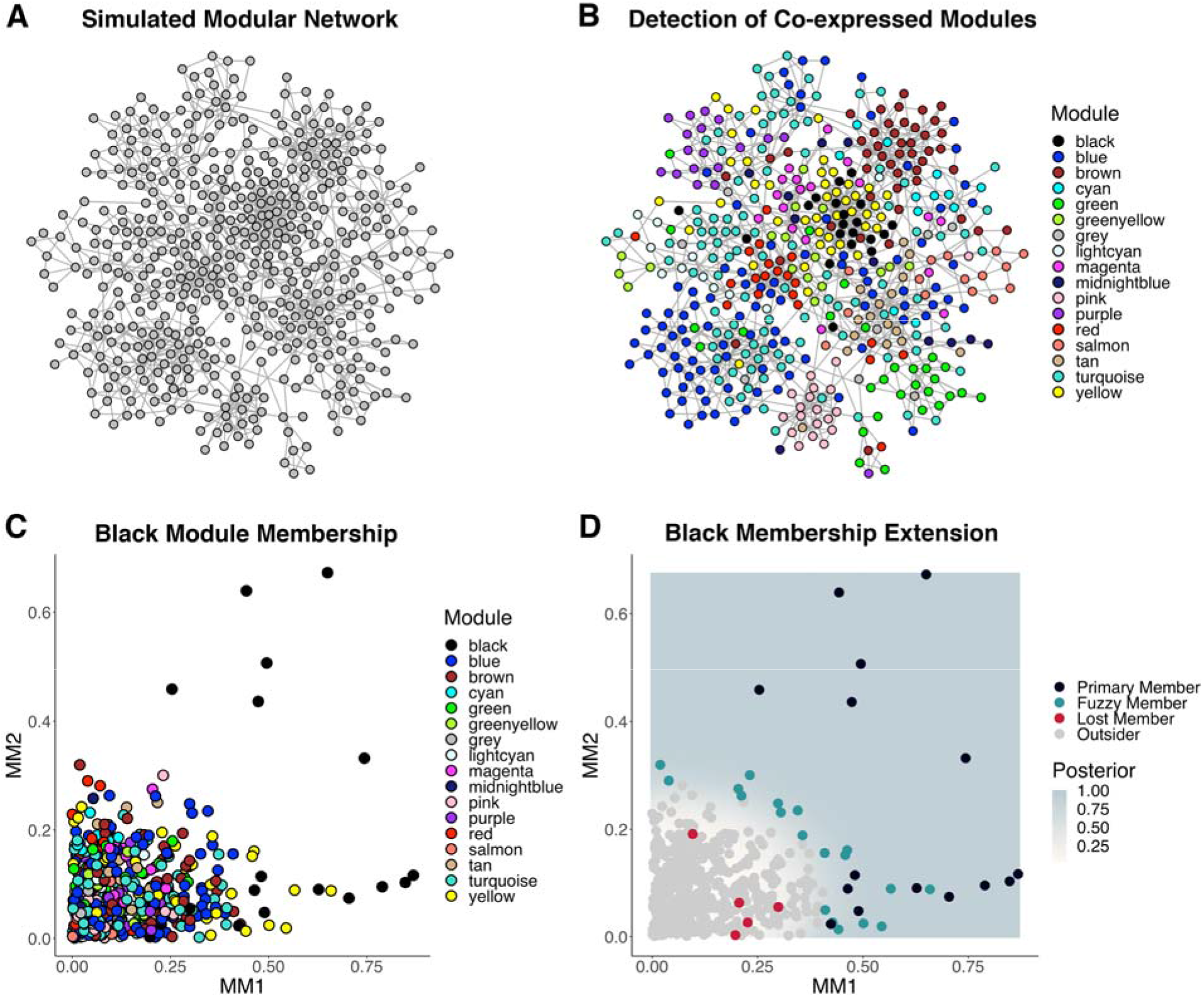
Module Detection and Extension. A) A simulated modular network with 500 nodes using the Lancichinetti–Fortunato–Radicchi benchmark. B) Detection of co-expression modules from simulated expression data where nodes are colored by their primary module. C) Nodes scored by their first (MM1) and second (MM2) membership in the black module. D) Nodes are assigned a membership probability *M_p_* (posterior) to the black module based on MM1/MM2. Black module membership is extended to non-member nodes with *M_p_* > 0.75 (Fuzzy Members).

### Extended Modules

We utilize a previously established concept in module detection called module membership (*MM*) to allow modules to contain overlapping sets of genes. Genes are assigned a membership score across all modules, where the membership of gene *i* in module *q* is the correlation of *i* and a module eigengene *E^(q)^*, which for the *q*^th^ module, is the first principal component of the expression profiles of genes within *q*, thus *MM* = / *bicor(x_i_, E^(q)^*) /. *MM* helps identify genes associated with multiple modules. Here we consider the *MM* of genes derived from the first *(MM1)* and second principal component *(MM2)*, which together often account for a majority of the variation of genes within a given module (Fig. 1C).

For each module, we consider non-member genes (outsiders) to become fuzzy members by comparing their first and second membership scores to primary member genes. This approach is formalized as a classification problem where we perform quadratic discrimant analysis (QDA) to classify genes (with some membership probability, *M_p_*) (Fig. 1D). In practice, control over the constraint levels is desirable. Thus, the probability of membership is a threshold that can be adjusted. After classification, modules are extended from their original members to the inclusion of fuzzy members. In this case, the graphical search only considers the interaction of genes *i* and *j* if they share one or more modules. Rather than constraining the entire graph, we employ a divide and conquer (DAQ) strategy for extended modules where we learn a graph for each module independently in parallel and then merge these sub-graphs into a final network. Conceptually with this modular approach, we are prioritizing local dependencies, by only accepting inferred edges if they are valid under all local conditions or subgraphs where they are tested.

We conducted simulation studies to compare methods for constraint-based structure learning on a single network with 300 nodes using the Lancichinetti–Fortunato–Radicchi (LFR) benchmark^16^. Simulated graph structures were used to simulate multivariate Gaussian data for an increasing number of samples ranging from 20 to 150, with the maximum sample size (*n* = 150) still significantly smaller than the data dimensions (*p* = 300). Overall, the constraint-methods performed better (by F1) than without constraints (Default) (Fig. 2A-C). The constrained-based methods managed to significantly reduce the number of false positives (FP) particularly with smaller sample sizes, which is advantageous when we are primarily interested in detecting high confidence interactions to generate hypotheses that can be validated in an experimental setting. Additionally, these methods maintained true positive (TP) levels similar to those achieved with an unconstrained search, thus resulting in overall improved performance by F1. While constraints based on isolated modules had the fewest FP, the resulting disconnected global graph structure makes downstream network analysis challenging. Thus, we prefer constraints based on extended modules. The efficiency and scalability of constraint-based structure learning with extended modules was further improved by using the DAQ strategy without a loss of overall performance.

**Figure 2.**
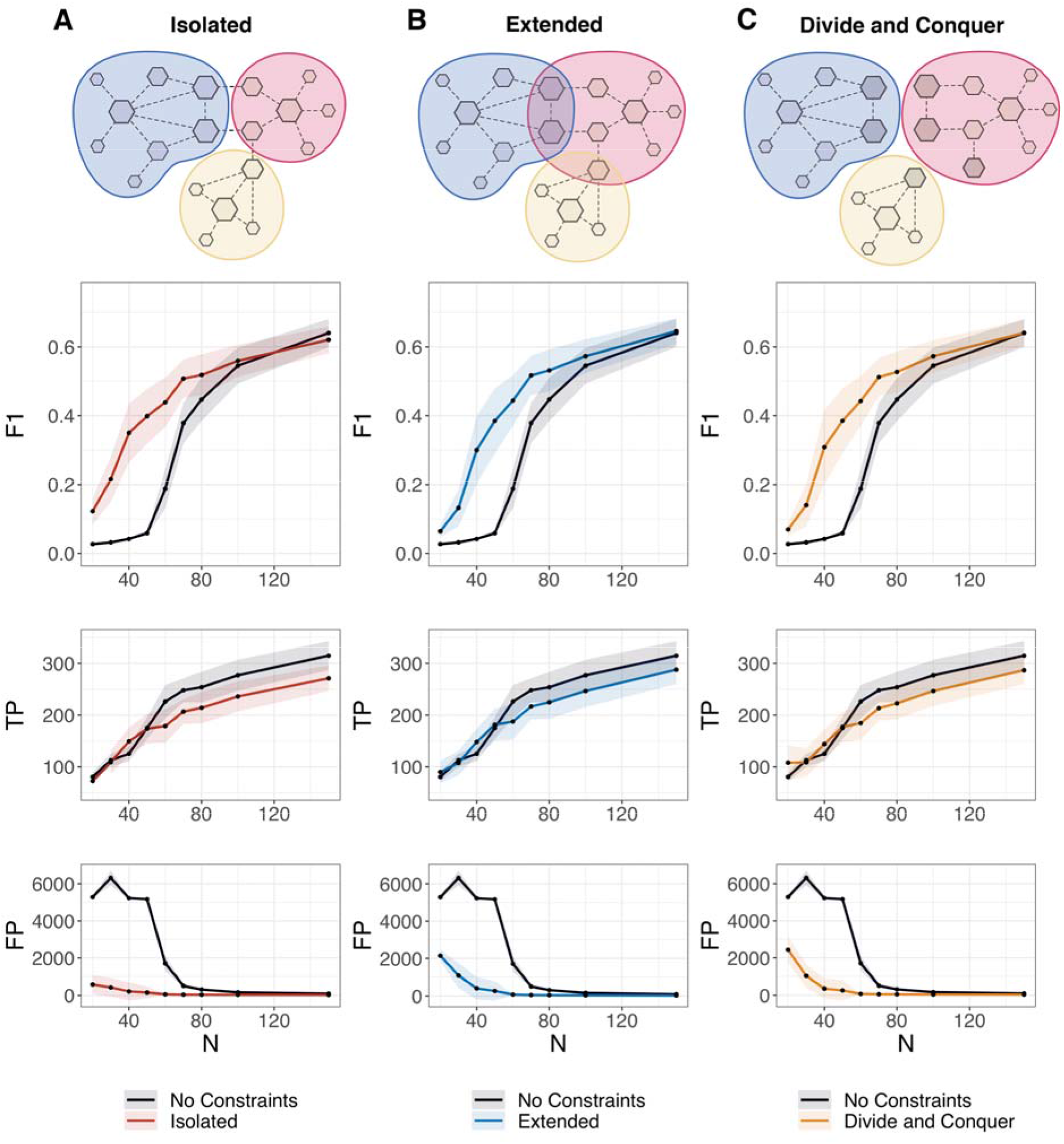
Simulation Studies for Constraint-based Methods. Performance of structure learning using constraint-based methods on a single network (*p* = 300) across an increasing sample range from 20 to 150 evaluated by the mean and standard deviation (*iterations* = 25) of F1 score (F1), true positives (TP), and false positives (FP). Isolated (A), Extended (B), and Divide and Conquer (C) approaches were compared to a baseline performance where no constraints were used.

### Shared Learning Increases the Equivalent Sample Size

When multiple networks are needed to model specializations of a common domain (e.g., different sub-types of breast cancer), encouraging shared learning of structural features is beneficial for each network individually. To implement this approach, we organize networks into a hierarchy, building a rooted tree of related networks. Leaf networks represent individual sample groups while networks higher in the hierarchy represent sample group supersets. Networks are inferred iteratively from the root down to the leaves, where at each level, the estimated posterior edges of the parent network are used to determine the prior distribution of edges in estimating the child network. The result is a hierarchy where internal and leaf networks represent shared and distinct network structures, respectively.

We simulated a hierarchy of similar networks to determine if shared learning further improves a constraint-based approach to network inference. To simulate network hierarchies, we employed an extension of the Barabási–Albert algorithm^17^ - using the concepts of growth and preferential attachment - where multiple networks were simulated from a shared seed graph (modeled via LFR), resulting in a hierarchy of graphs with a known edge similarity (Fig. 3A-B). With sample sizes at the lower extreme, we found the use of a prior network to be highly beneficial in addition to the incorporation of constraints (Fig. 3C). Conversely, as *n* approached *p*, inference became tractable without constraints or prior information, shared learning limited the ability to detect new TP, and we saw diminishing returns in F1. This suggests that these methods are most applicable when sample size is small relative to the desired feature set, which is highly relevant in the context of biological network inference where n ≪ p. We refer to this use of constraint-based shared learning as *SHINE* (**S**tructure Learning for **Hi**erarchical **Ne**tworks).

**Figure 3.**
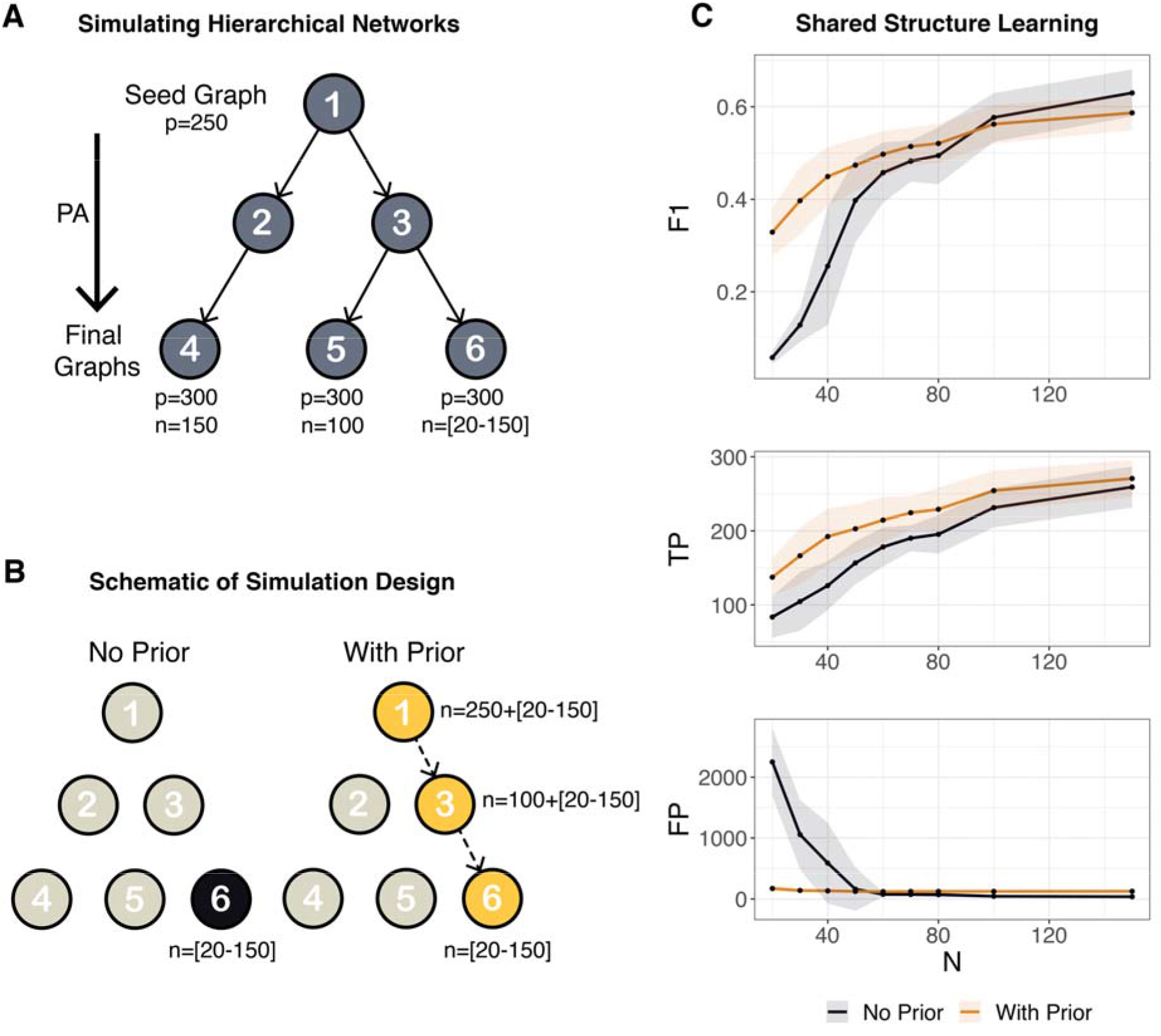
Simulation Studies for Shared Learning on Hierarchical Networks. A) A schematic of generating a network hierarchy by simulating independently diverging graph structures with a known edge similarity, arriving at final graph structures use to simulate data for sample groups 4, 5, and 6. B) A schematic of the simulation design whereby shared learning of network 6 (using samples from group 6) incorporates prior information from network 3 (pooled samples from groups 5-6) which incorporates prior information of network 1 (pooled samples from groups 4-6). C) Performance of structure learning using the divide and conquer constraint-based method with and without prior information on a network 6 across an increasing group 6 sample range from 20 to 150 evaluated by the mean and standard deviation (*iterations* = 25) of F1 score (F1), true positives (TP), and false positives (FP).

### Comparing SHINE with Other Methods

We compared *SHINE* to other available network inference methods in a network hierarchy context. In particular, we compared the accuracy of these methods in recovering the gold leaf network of the hierarchy in Fig. 3B. We used representatives of three popular categories including correlation-based, mutual information-based, and tree-based approaches. These include Weighted Correlation Network Analysis (WGCNA), Algorithm for the Reconstruction of Accurate Cellular Networks (ARACNe), and Gene Network Inference with Ensemble of Trees (GENIE3)^18,19^. WGCNA and ARACNe find potential gene interactions by measuring the correlation and mutual information between gene pairs, respectively. Additionally, ARACNe uses the data processing inequality (DPI) to remove the weakest edge from every fully connected node triplet. GENIE3 infers interactions by solving a regression problem for each gene. These methods generate a ranked list of weighted interactions which can be used to construct a gene regulatory network by choosing an interaction threshold. *SHINE* had a higher overall performance compared to these methods, with higher F1 scores across increasing sample sizes. *SHINE* had larger F1 improvement for small sample sizes (relative to the number of nodes) (Table 1). Furthermore, it should be emphasized that the optimal threshold of the compared methods is not a-priori known. In contrast, *SHINE’s* adjacency matrices were constructed by a-priori setting a probabilistic threshold for edges (*p = 0.9*), with the results largely insensitive to the threshold choice (*p = 0.85-0.95*). This is preferred in real application settings, where the correct threshold is in general unknown.

**Table 1.** The average F1-score of learned networks with increasing sample size. SHINE was compared to three methods, each using various cut points for retaining the top percentage of predicted edges.

| Method | Cut | n=20 | n=30 | n=40 | n=50 | n=60 | n=70 | n=80 | n=100 | n=150 |
| --- | --- | --- | --- | --- | --- | --- | --- | --- | --- | --- |
| SHINE | NA | 0.329 | 0.397 | 0.449 | 0.474 | 0.497 | 0.514 | 0.521 | 0.562 | 0.587 |
| GENIE3 | 0.1% | 0.097 | 0.126 | 0.135 | 0.147 | 0.149 | 0.149 | 0.152 | 0.150 | 0.154 |
| GENIE3 | 1.0% | 0.222 | 0.304 | 0.355 | 0.391 | 0.413 | 0.436 | 0.444 | 0.469 | 0.505 |
| GENIE3 | 2.5% | 0.191 | 0.247 | 0.289 | 0.307 | 0.322 | 0.339 | 0.342 | 0.369 | 0.392 |
| ARACNe | 0.1% | 0.121 | 0.137 | 0.147 | 0.153 | 0.152 | 0.156 | 0.157 | 0.156 | 0.156 |
| ARACNe | 1.0% | 0.226 | 0.303 | 0.365 | 0.397 | 0.430 | 0.448 | 0.464 | 0.489 | 0.549 |
| ARACNe | 2.5% | 0.027 | 0.090 | 0.108 | 0.136 | 0.148 | 0.168 | 0.195 | 0.165 | 0.198 |
| WGCNA | 0.1% | 0.122 | 0.127 | 0.130 | 0.136 | 0.138 | 0.139 | 0.145 | 0.142 | 0.141 |
| WGCNA | 1.0% | 0.259 | 0.314 | 0.336 | 0.376 | 0.387 | 0.398 | 0.409 | 0.414 | 0.437 |
| WGCNA | 2.5% | 0.215 | 0.257 | 0.283 | 0.307 | 0.322 | 0.338 | 0.339 | 0.353 | 0.371 |

### Evaluation on Pan-Cancer Data

We reconstructed Pan-Cancer networks from The Cancer Genome Atlas (TCGA) RNA-seq data, to evaluate *SHINE* and demonstrate its scalability. Cancer types were organized into a four-level hierarchy, starting with all primary tumors (L0), then further stratified into organ system (L1), tumor type (L2), and tumor subtype (L3) (Supplementary Table 1)^20^. Due to our focus on cancer regulatory networks we learned networks from 1157 genes involved in 14 major pathways encompassing hallmark capabilities of cancer (Supplementary Table 2)^21^. The rationale being that detection of changes in connectivity across such networks will likely represent differences in the outcome of cellular processes underlying tumor initiation and progression. We built extended modules on L0 and shared these constraints to learn L1 networks using the DAQ approach. L2 networks were learned using L1 as a prior based on the pre-defined hierarchy (Supplementary Fig. 1).

We found networks largely clustered according to their prior network (e.g., L2 networks that shared an L1 network as a prior were more similar) (Fig. 4A). We compared learned and randomly simulated networks (via *Barabási–Albert*^17^ and *Erdős–Rényi*^22^ models) to experimentally derived protein-protein interactions (PPI)^23^. Inferred interactions in each of the cancer networks were far more significantly enriched for PPIs than interactions in the random networks (BA and ER). Furthermore, learned networks shared similar graph properties with the PPI network, such as low density (sparsity) and high global and local clustering coefficients (transitivity and clustering respectively), which are key properties of real biological networks (Table 2)^9^.

**Figure 4.**
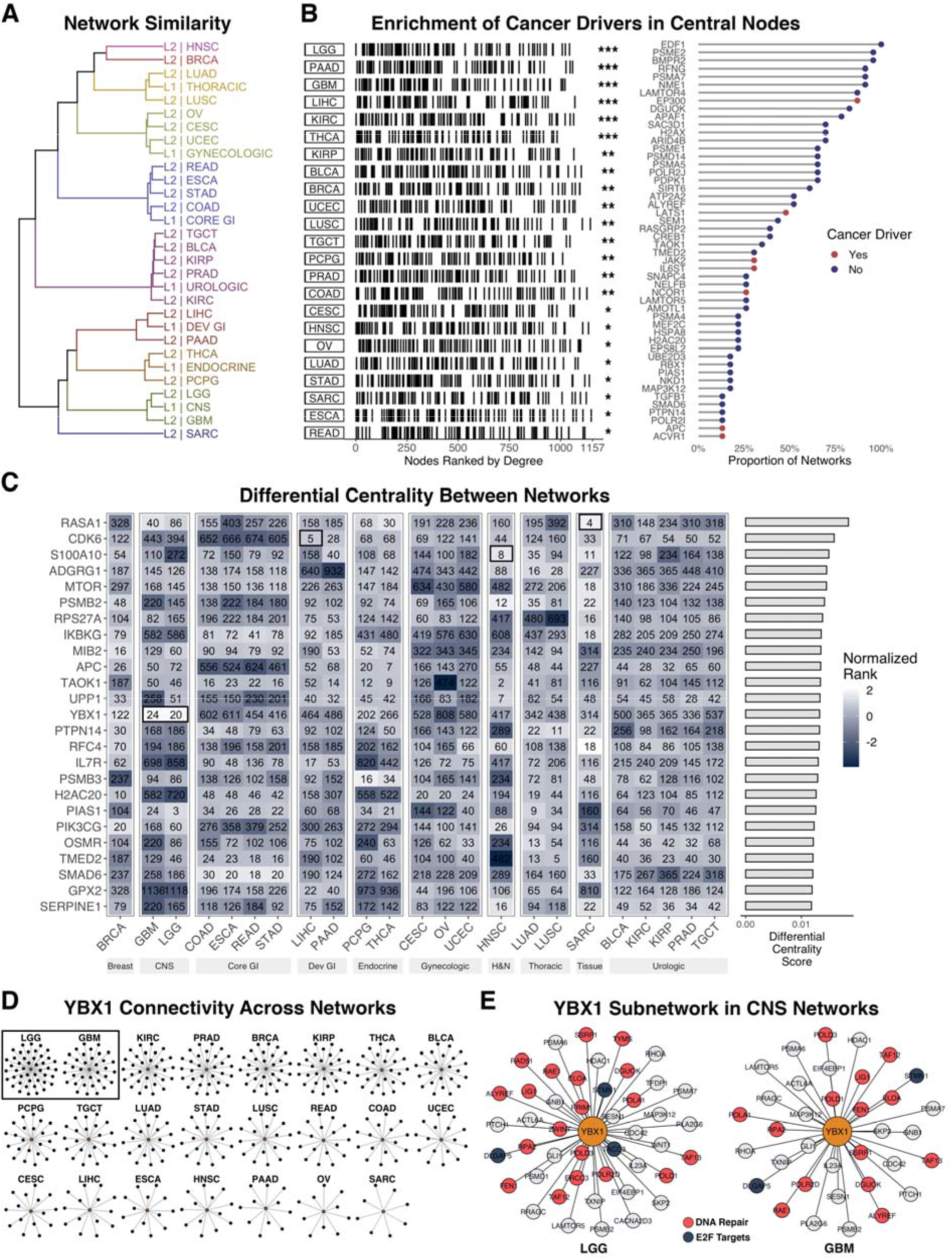
Pan-Cancer Networks. A) Network similarity clustered by Kendall rank correlation of degree centrality, where L1/L2 signify the network level in the pre-defined hierarchy. B) Enrichment of previously identified cancer driver genes in networks where nodes are ranked by degree (*left*). High-degree nodes (top 25) in at least one network. The top 50 most frequently present genes across networks are shown (*right*). C). Top 25 differentially central genes (by degree) between networks ranked by their differential centrality score. D) YBX1 (*orange*) and its first-degree connections in each network. E) YBX1 subnetwork of first-degree connections in CNS networks (LGG and GBM).

**Table 2.** Common graph-level and node-level measures computed for learned Pan-Cancer networks. Inferred network edges were tested against a database of experimentally validated PPIs in human and compared with randomly simulated networks.

| Network | Node | Edges | Density | Transitivity | Clustering | Assortivity | Overlap | P-Value | FDR |
| --- | --- | --- | --- | --- | --- | --- | --- | --- | --- |
| KIRC | 1,157 | 14,868 | 0.022 | 0.125 | 0.170 | 0.115 | 638 | 4.8 x 10<sup>(-87)</sup> | 1.2 x 10<sup>(-85)</sup> |
| KIRP | 1,157 | 13,664 | 0.020 | 0.118 | 0.160 | 0.147 | 602 | 2.2 x 10<sup>(-86)</sup> | 2.8 x 10<sup>(-85)</sup> |
| STAD | 1,157 | 10,036 | 0.015 | 0.096 | 0.128 | 0.129 | 488 | 1.8 x 10<sup>(-83)</sup> | 1.5 x 10<sup>(-82)</sup> |
| PRAD | 1,157 | 14,163 | 0.021 | 0.120 | 0.167 | 0.121 | 604 | 2.2 x 10<sup>(-81)</sup> | 1.4 x 10<sup>(-80)</sup> |
| BLCA | 1,157 | 13,453 | 0.020 | 0.115 | 0.156 | 0.139 | 582 | 1.2 x 10<sup>(-80)</sup> | 6.1 x 10<sup>(-80)</sup> |
| TGCT | 1,157 | 13,119 | 0.020 | 0.113 | 0.155 | 0.132 | 567 | 2.1 x 10<sup>(-78)</sup> | 8.7 x 10<sup>(-78)</sup> |
| READ | 1,157 | 9,183 | 0.014 | 0.093 | 0.123 | 0.117 | 448 | 4.4 x 10<sup>(-77)</sup> | 1.6 x 10<sup>(-76)</sup> |
| ESCA | 1,157 | 9,048 | 0.014 | 0.092 | 0.125 | 0.143 | 443 | 1.1 x 10<sup>(-76)</sup> | 3.4 x 10<sup>(-76)</sup> |
| COAD | 1,157 | 10,790 | 0.016 | 0.101 | 0.137 | 0.113 | 491 | 8.6 x 10<sup>(-75)</sup> | 2.4 x 10<sup>(-74)</sup> |
| THCA | 1,157 | 10,977 | 0.016 | 0.097 | 0.128 | 0.103 | 480 | 6.5 x 10<sup>(-68)</sup> | 1.6 x 10<sup>(-67)</sup> |
| LUAD | 1,157 | 9,998 | 0.015 | 0.094 | 0.128 | 0.066 | 448 | 2.2 x 10<sup>(-66)</sup> | 5.0 x 10<sup>(-66)</sup> |
| LIHC | 1,157 | 7,449 | 0.011 | 0.077 | 0.104 | 0.118 | 370 | 1.0 x 10<sup>(-65)</sup> | 2.1 x 10<sup>(-65)</sup> |
| LGG | 1,157 | 10,557 | 0.016 | 0.100 | 0.129 | 0.115 | 456 | 4.8 x 10<sup>(-63)</sup> | 9.2 x 10<sup>(-63)</sup> |
| PAAD | 1,157 | 6,119 | 0.009 | 0.071 | 0.093 | 0.097 | 322 | 8.2 x 10<sup>(-63)</sup> | 1.5 x 10<sup>(-62)</sup> |
| LUSC | 1,157 | 9,489 | 0.014 | 0.092 | 0.125 | 0.078 | 423 | 4.0 x 10<sup>(-62)</sup> | 6.6 x 10<sup>(-62)</sup> |
| UCEC | 1,157 | 10,016 | 0.015 | 0.098 | 0.132 | 0.114 | 436 | 3.2 x 10<sup>(-61)</sup> | 5.1 x 10<sup>(-61)</sup> |
| GBM | 1,157 | 8,275 | 0.012 | 0.086 | 0.117 | 0.136 | 384 | 1.0 x 10<sup>(-60)</sup> | 1.5 x 10<sup>(-60)</sup> |
| PCPG | 1,157 | 8,607 | 0.013 | 0.084 | 0.108 | 0.111 | 393 | 2.3 x 10<sup>(-60)</sup> | 3.2 x 10<sup>(-60)</sup> |
| OV | 1,157 | 8,855 | 0.013 | 0.091 | 0.123 | 0.125 | 400 | 3.1 x 10<sup>(-60)</sup> | 4.0 x 10<sup>(-60)</sup> |
| CESC | 1,157 | 8,773 | 0.013 | 0.092 | 0.124 | 0.126 | 396 | 1.5 x 10<sup>(-59)</sup> | 1.8 x 10<sup>(-59)</sup> |
| BRCA | 1,157 | 10,350 | 0.015 | 0.099 | 0.134 | 0.097 | 434 | 1.2 x 10<sup>(-56)</sup> | 1.4 x 10<sup>(-56)</sup> |
| HNSC | 1,157 | 6,578 | 0.010 | 0.078 | 0.109 | 0.085 | 309 | 3.9 x 10<sup>(-50)</sup> | 4.4 x 10<sup>(-50)</sup> |
| SARC | 1,157 | 3,639 | 0.005 | 0.061 | 0.081 | 0.049 | 172 | 7.2 x 10<sup>(-29)</sup> | 7.8 x 10<sup>(-29)</sup> |
| BA<sup>1</sup> | 1,157 | 1,156 | 0.002 | 0.000 | 0.000 | -0.116 | 30 | 3.2 x 10<sup>(-2)</sup> | 3.3 x 10<sup>(-2)</sup> |
| ER<sup>1</sup> | 1,157 | 9,917 | 0.015 | 0.015 | 0.015 | -0.002 | 179 | 5.0 x 10<sup>(-1)</sup> | 5.0 x 10<sup>(-1)</sup> |
| PPI<sup>2</sup> | 1,157 | 12,021 | 0.018 | 0.196 | 0.277 | -0.054 | NA | NA | NA |

<sup>1</sup>Simulated Networks
<sup>2</sup>Experimentally Validated

The degree distributions across networks were largely scale free, and highly connected nodes were enriched for known cancer drivers in all networks (*p = 4.44e-16*) (Fig. 4B left). Highly connected nodes were conserved across a majority of the networks; many of which are implicated in the initiation, progression, and invasiveness of tumors. For example, EDF1 was a top hub in every network and PSME2^24^, BMPR2^25^, RFNG, PSMA7^26^, NME1^27^, LAMTOR4^28^, and EP300^29^ were top hubs in at least 80% of networks. Interestingly, while cancer drivers tend to be more central across networks, the most connected hubs are not necessarily cancer drivers (Fig. 4B right).

To identify tumor type-specific differences, we scored nodes by their differential degree-centrality using a normalized rank score across networks. Many of the nodes found to be differentially central to a particular network(s) were previously implicated in their respective tumor type (Fig. 4C). Using the top three as examples: RASA1 was shown to be involved in the development of some sarcomas^30,31^, CDK signaling was found to be a potential therapeutic target in hepatocellular carcinoma^32^, and S100A4 was shown to maintain cancer-initiating cells in head and neck cancers^33^. Interestingly, YBX1 - a transcription factor involved in DNA repair, highly expressed in cancers, and known to promote cell growth and to inhibit differentiation in glial tumors - is highly central in both CNS subtype networks (LGG and GBM) (Fig. 4D)^34^. Furthermore, in these tumor types, YBX1 subnetworks are enriched for DNA repair genes (*p* = *5.8e-07* and *p* = *4.3e-05*; LGG and GBM respectively) and connected to multiple E2F targets (Fig. 4E).

### Learning Networks for Breast Cancer Subtypes

We applied *SHINE* to the analysis of TCGA Breast Cancer RNA-seq data to learn subtype-specific networks corresponding to the PAM50 molecular classification into Luminal A, Luminal B, HER2-enriched, and Basal-like (the Normal-like subtype was excluded because of too small a sample size, *n* = 40)^35^. To further boost the equivalent sample size, we used experimentally validated PPIs from breast cancer-related cell lines (e.g., MCF-7 and MDA-MB-231) as a structural prior or starting graph in the network hierarchy (Supplementary Fig. 2)^36^. We found networks had similar community structure, and many of the top genes ranked by centrality measures were conserved across subtype networks (Fig. 5B *top*). Interestingly, Luminal-B and HER2-enriched networks were most similar while Luminal-A was most dissimilar when comparing networks by centrality rankings.

**Figure 5.**
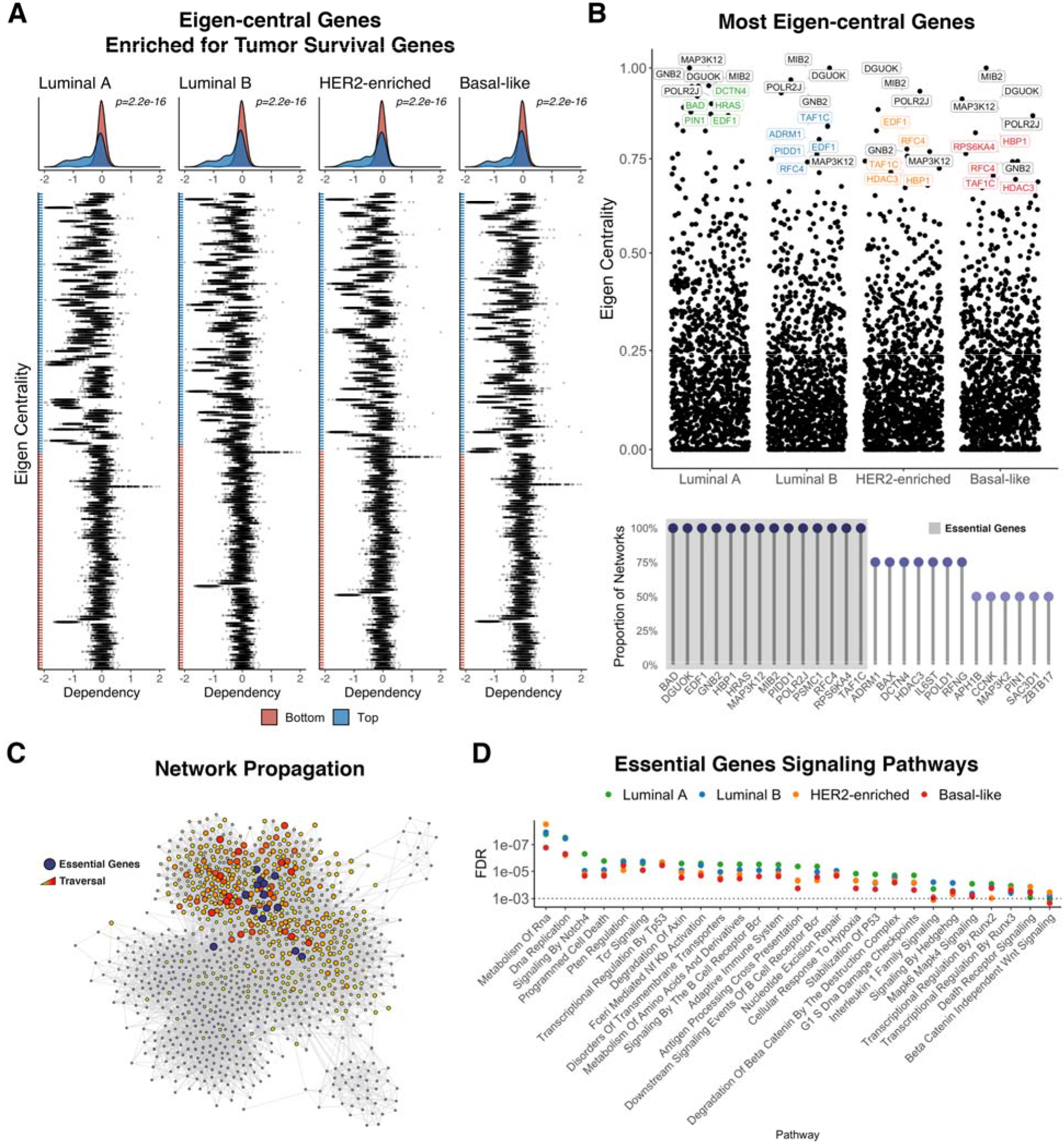
Breast Cancer Essential Genes. A) Genes within the top and bottom 100 eigen-centrality ranking across breast cancer networks scored by their genetic dependency across various cell lines via the Cancer Dependency Map Project (DepMap). A negative dependency value indicates higher essentiality to tumor survival. Genes not present in the DepMap are omitted. B) Top 10 eigen-central genes across networks (*top*). Top 25 eigen-central genes across multiple networks. Highlighted are 14 genes identified as highly central in all networks (*bottom*). C) A visualization of network propagation using the 14 eigen-central breast cancer genes as seed nodes where a red node indicates high traversal and close proximity to the seed genes. D) Biological pathway enrichment of highly traversed genes.

We found eigen-central genes in the networks to be enriched for genes known to be essential for tumor survival identified by the Dependency Map Project (Fig. 5A)^37^. A highly central node indicates high network influence; thus, we expect highly eigen-central genes across inferred networks to play a central regulatory role in maintaining breast tumorigenesis and survival. Among these are 14 genes - BAD, DGUOK, EDF1, GNB2, HBP1, HRAS, MAP3K12, MIB2, PIDD1, POLR2J, PSMC1, RFC4, RPS6KA4, TAF1C - ranked within the top 25 by eigen-centrality in all four subtypes (Fig. 5B *bottom*). Their highly conserved rank suggests they are likely to be central players driving and maintaining breast cancer. Interestingly, these central players localize in close proximity to each other in all network subtypes (Fig. 5C). Network propagation seeded with these essential genes, and subsequent pathway enrichment analysis on highly traversed nodes supports the hypothesis that these genes facilitate crosstalk between DNA maintenance/repair and immune-related pathways (Fig. 5D).

We found a community identified via Walktrap^38^ community detection enriched for known breast cancer disease genes in Luminal A (*FDR* = *0.00035*) and Basal-like (*FDR* = *0.0026*) networks (Fig. 6A)^39^. This disease neighborhood was present in both subtypes, containing the same set of 31 disease genes and enriched for immune-related signaling pathways (Fig. 6B). We used the guilt-by-association principle to predict novel breast cancer genes by performing network propagation, seeded with the existing disease genes (Fig. 6C-D)^40,41^. It has been postulated that drugs may be more effective through network-based actions to target multiple disease genes^42^. Of note, not only is PDGFRB within close proximity to all disease genes in both networks, but it is also the most connected and eigen-central hub of the neighborhood. PDGFRB may be a prime candidate for disease modulation and is listed as a targetable gene (via Cediranib, Masitinib, Pazopanib) from the Genomics of Drug Sensitivity in Cancer^43^. Furthermore, PDGF signaling has been found to play an active role in breast tumor progression and PDGFRB has previously been theorized - along with its cognate ligand PDGFB - to be a potential therapeutic target for multiple human cancers, including breast^44^. Interestingly, PDGFRB/PDGFB are in closer proximity to disease genes in the Basal-like subnetwork further supporting evidence for its role in tumor aggression^45^.

**Figure 6.**
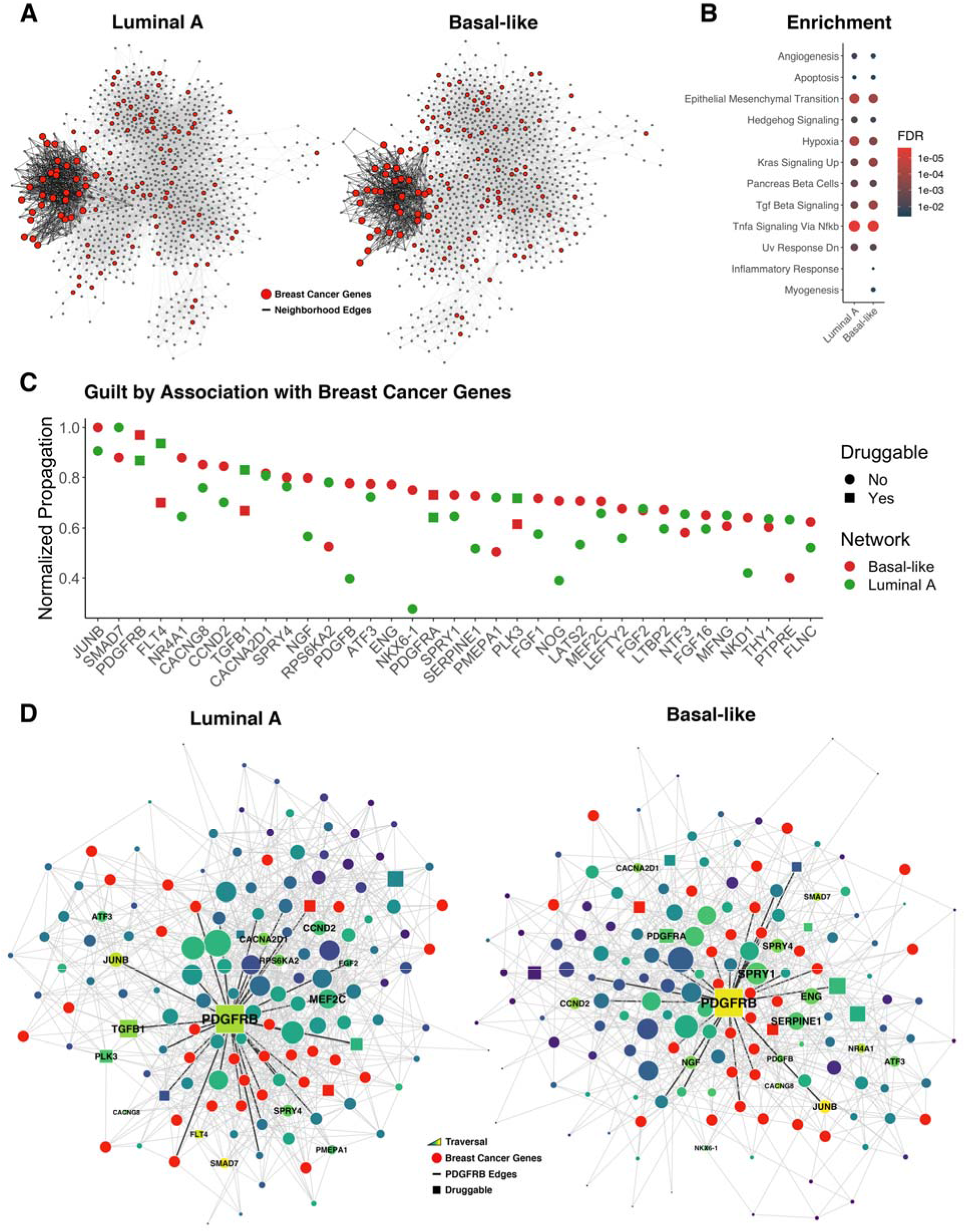
Breast Cancer Disease Neighborhood. A) A single community detected in breast cancer subtype networks enriched for known breast cancer disease genes. B) Pathway enrichment of the identified disease neighborhoods. C) Proximity of neighborhood genes to known breast cancer disease genes identified through network propagation. D) Subnetwork visualization of disease neighborhoods in Luminal A and Basal-like networks.

## DISCUSSION

In this report we present a novel multi-pronged approach called *SHINE* for learning biological Markov networks from limited sample sizes. We exploit known organizing principles of biological networks to limit the model parameters of structure learning and encourage shared learning of multiple networks to boost the equivalent sample size. This approach reduces the complexity of the search space, allows related networks to share data, and takes advantage of a-priori structural information, resulting in higher overall performance with fewer false positives. There is a cost of slightly more false negatives, however this tradeoff is advantageous in the context of biological network inference where n≪p, since we are primarily interested in predicting high confidence interactions for hypothesis generation. We apply *SHINE* to reconstruct tumor-specific networks from TCGA data as well as a focused analysis on breast cancer and find inferred networks exhibit expected graph properties of real biological networks, recapture previously validated interactions, and recapitulate findings in literature.

We found *SHINE* to be highly beneficial for learning networks, however this approach is only suitable when co-expression modules can be detected from the data. In simulation studies, we found that at very low samples sizes (*n* < 20 and *p* = 300), a failure to build appropriate constraints led to poor inference performance. While *SHINE* is largely data-driven and there are few parameters to optimize, it is useful to adjust the level of fuzziness in defining extended modules, which can be controlled via the threshold or decision boundary for secondary membership M_*p*_. M_*p*_ can vary between unconstrained (M_*p*_ = 0) to isolated (M_*p*_ = 1), providing a level of control over high and low tolerance of false positives respectively. There is evidence that interactions across modules are sparse, while intra-modular connections are dense, suggesting that a good rule of thumb is to favor strong constraints with a higher M_*p*_ threshold^46^. Furthermore, *SHINE* could be extended to include more than the first two eigengenes to account for more variation in the calculation of membership or take advantage of a wide variety of alternative methods for module detection, some of which detect overlapping modules by default^11^.

The learning process is rendered computationally highly efficient by the parallelization made possible via the DAQ approach, as well as by the simultaneous inference of independent networks in a given hierarchy. By using a DAQ approach, the upper bound of *p* scales with the number of sufficiently small modules used as constraints. In our analysis, we aimed for extended modules smaller than 300 genes, thus using an *M_p_* = 0.9. We believe this is a reasonable limit as over 97% of biological pathways defined in Kegg, Reactome, and Biocarta are smaller than 300 genes. Therefore, *SHINE* scales well for datasets where a large number of small and tightly clustered co-expression modules can be identified. Parallelization was fully automated through adoption of Nextflow to manage network workflows. Because some networks are dependent on others (e.g., a network is used as a prior in learning another), we can take advantage of the reactive properties of Nextflow – a language for defining portable, scalable, and reproducible data workflows – which solves and handles the process dependencies between parent and child networks. For example, the application to the Pan-Cancer hierarchy of networks required 656 individual jobs, many of which could be run in parallel, reducing our sequential run time from 52 to 4 hours.

The network hierarchy used to guide shared learning can be specified in advance if it is known, or it can be learned from the data, or a hybrid approach where both “expert knowledge” and a data driven approach to learning can be combined. The use of prior information is highly flexible in that one could incorporate only network constraints, previously learned networks (as a prior distribution of edges), existing network structures (as a seed graph), or a combination. Last, while we focus on human gene regulatory networks, it is reasonable to extend this approach to other biological systems and data types, both bulk and single cell data, as well as to other high-throughput omics layers such as proteomics or metabolomics, among others.

## METHODS

### Module Detection and Extension

Genes are clustered by their co-expression similarity *s_ij_* – measured by the absolute value of the biweight midcorrelation coefficient: *s_i,j_* = |*bicor*(*x_i_, x_j_*) | as well as a soft thresholding value *ß* which pushes spurious correlations to zero, resulting in a symmetric *p x p* weighted adjacency matrix 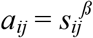. Co-expression modules are detected using hierarchical clustering of a topological overlap dissimilarity transformation *d_i,j_* of *a_ij_* resulting in *Q* modules. Genes are assigned a membership score across all modules, where the membership of gene *i* in module *q* is the correlation of *i* and a module eigengene *E^(q)^*, which for the *q*^th^ module, is the first principal component of the expression profiles of genes within *q*, thus *MM* = | *bicor*(*x_i_, E^(q)^*) /. Within each module, each gene is assigned a probability of membership through quadratic discrimant analysis (QDA) based on MM1/MM2. Module membership is extended to non-member genes above a membership probability *M_p_*.

### Simulation Studies

Single network simulations were performed using undirected graphs (*p = 300*) randomly generated using the Lancichinetti–Fortunato–Radicchi (LFR) benchmark (*tau1 = 3, tau2 = 2, mu = 0.08*). The LFR model generates graphs with overlapping community structures, where both degree and community size follow a power law. The graph structures were used to generate multivariate Gaussian data using the R package *BDgraph*^3^ for samples ranging from 20 to 150. Structure learning with constraint-based approaches including Isolated, Extended, and Divide and Conquer (DAQ) was performed for each sample size - repeated 25 times - where a posterior edge probability above 0.9 was considered an edge in the final network. Networks were evaluated based on an F1-score = 2TP/(2TP+FP+FN’) where TP, FP, and FN are the number of true positives, false positives, and false negatives respectively and accounts for a balance of detecting TP while limiting FP. Simulations with multiple networks were performed under similar conditions by generating a hierarchy of networks with a known edge similarity. This was done by using an initial graph (*p = 250*) generated by LFR (*tau1 = 3, tau2 = 2, mu = 0.08*) to simulate independently diverging network structures (via the Barabási–Albert procedure for network growth and preferential attachment) - where divergent networks grow by an additional 25 nodes - arriving at three final graphs (*p = 300*) with an average 84% edge similarity calculated by the Szymkiewicz–Simpson coefficient^47^ of edges (Fig. 3A). The graph structures were used to generate multivariate Gaussian data using the R package *BDgraph*. Networks were estimated and evaluated with the DAQ constraint method with and without the use of prior network information using previously described methods across increasing samples ranging from 20 to 150 - repeated 25 times.

### Data Pre-Processing

The Cancer Genome Atlas (TCGA) RNA-Seq count matrices (generated with STAR 2-Pass and HTSeq - Counts) and metadata for all available tumor types was downloaded through the Genomic Data Commons (GDC) *gdc-client*^48^. We performed a variance-stabilizing transformation of the data using the R package *DESeq2* followed by a log-transformation^49^. We removed datasets with fewer than 150 primary tumors, resulting in the following 23 tumor types (L2) and 11 categories (L1): Breast - *BRCA* (*n* = 1102); Thoracic - *LUSC* (*n* = 502), *LUAD* (*n* = 533); Endocrine - *THCA* (*n* = 502), *PCPG* (*n* = 178); Head and Neck - *HNSC* (*n* = 500); CNS - *LGG* (*n* = 511), *GBM*(*n* = 156); Soft Tissue - *SARC* (*n* = 259); Gynecologic - *UCEC* (*n* = 551), *OV* (*n* = 374), *CESC* (*n* = 304); Urologic - *KIRC* (*n* = 538), *PRAD* (*n* = 498), *BLCA* (*n* = 414), *KIRP* (*n* = 288), *TGCT*(*n* = 150); Developmental GI - *LIHC* (*n* = 371), *PAAD* (*n* = 177); Core GI - *COAD* (*n* = 478), *STAD* (*n* = 375), *READ* (*n* = 166), *ESCA* (*n* = 161) (Supplementary Table 1). For the Pan-Cancer analysis, gene-wise expression levels were modified by taking the residuals after adjusting for tumor type.

### Gene Filtering

Networks were built on genes from the following major cancer pathways sourced from the Molecular Signatures Database (MSigDB)^50^ (v7.2.1) and Pathway Commons^51^ (v12.0) resulting in a unique set of 1160 genes, 3 of which were not available in the data, further reducing to 1157 final genes. The pathways and geneset sizes include MAPK Signaling (267), Notch Signaling (247), P53 Signaling (200), Jak Stat Signaling (155), DNA Repair (150), TGF-Beta Signaling (54), Oxidative Stress Response (45), PI3K Pathway (43), Wnt/Beta-Catenin Signaling (42), Mtor Signaling (40), Hedge Hog Signaling (36), Myc Signaling (25), RAS Signaling (24), Hippo Signaling (20), Cell Cycle (14) (Supplementary Table 2).

### Inference of Pan-Cancer Networks

We computed shared constraints derived from all primary tumors through the R package *shine* which extends methods implemented in the R package *WGCNA*^12^. The soft-thresholding power *2* was set to 3 (the lowest value from 1-20 for which the scale-free topology fit had at least an *R^2^ > 0.80*). Co-expression similarity was calculated using biweight midcorrelation and unsigned options. Hierarchical clustering was performed using the average agglomeration method. Adaptive branch pruning was performed using the hybrid method with deep split set to 4 and a minimum cluster size of 10. Lastly, the first module – which represents genes that fail to cluster into a co-expression module of a minimum size – was removed from downstream methods. The resulting 18 modules had sizes ranging from 13-269 with a mean of 63.89 and median of 33. Modules were extended with an *M_p_* = 0.9 extending their sizes to a range of 27-290 with a mean of 82.61 and median of 54. Networks were inferred using the structure learning R package *BDgraph*. We used the undirected Gaussian graphical model search method based on marginal pseudo-likelihood using the birth-death Markov chain Monte Carlo algorithm. Learning constraints were applied with the previously described DAQ strategy. When learning a child network from a parent network, the posterior distribution of edges of the parent is used as the prior in learning the child.

### Inference of Breast Cancer Networks

We computed shared constraints as previously described on all primary breast tumors (*n* = 1102) - DESeq2-log normalized and unadjusted for tumor type - from the following subtype classifications: Luminal A (*n* = 564), Luminal B (*n* = 215), HER2-enriched (*n* = 82), and Basal-like (*n* = 189), Normal-like (*n* = 40), and Indeterminate (*n* = 12). We did not learn an individual network for Normal-like due to its small sample size. The resulting 24 modules had sizes ranging from 11-205 with a mean of 44.38 and median of 28.50. Modules were extended with an *M_p_* = 0.9 extending their sizes to a range of 23-233 with a mean of 68.17 and median of 51.50. Additionally, we used experimentally validated PPIs from MCF-7, MDA-MB-231, MCF-10A, MCF-10AT cell lines annotated in *Federico et. al.* as a structural prior or starting graph in the network hierarchy^36^. Networks were then inferred using the previous described methods.

### Hierarchical Workflows

The heavy computational demand for high-dimensional graphical modeling requires networks to be learned in parallel on high performance computing platforms. Because lower-level networks depend on higher-level networks in the hierarchy, we employed the reactive workflow scripting language Nextflow^52^ (DSL2) to manage workflows given a network hierarchy within a containerized computing environment via Docker. Together, this enables automatic job parallelization, failure recovery, and portability to various cloud/cluster architectures, resulting in reproducible workflows that are easy to manage. We have developed a collection of Nextflow DSL2 modules for running *SHINE* as well as a small command line utility written in Python for dynamically generating these workflows based on the hierarchical structure of multiple networks.

### Calculation of Network Properties and Similarity

Networks were formatted into graph objects using the R package *igraph*^53^. We used built-in functions to compute common graph-level and node-level measures such as density, transitivity (global clustering coefficient), clustering (local clustering coefficient), and assortivity. We compared inferred interactions to human experimentally derived protein-protein interactions (PPI) from the Human Integrated Protein-Protein Interaction Reference (HIPPIE)^23^ v2.2. Significance of intersecting sets was computed using a Fischer’s exact test. We also compared the resulting network properties to 1000 iterations (taking the median values) of randomly simulated undirected networks built with the Erdős–Rényi (ER) and Barabási–Albert (BA) models. We computed network similarity or distance via the Kendall rank correlation of nodes ranked by degree for all pairwise networks. Hierarchical clustering was then performed on this distance matrix and visualized as a dendrogram.

### Enrichment of Cancer Drivers in Central Nodes

Nodes in each network were ranked by their degree-centrality and tested for enrichment of cancer driver genes identified in *Bailey et. al*. in their TCGA Pan-Cancer analysis by a Kolmogorov-Smirnov test^54^. The overall significance was computed by comparing the distribution of p-values arising from each enrichment test to a uniform distribution.

### Computing Differential Centrality Scores

Nodes were ranked by a centrality measure and compared between networks to identify genes differentially connected or central in one or more networks. For *n* nodes, we represent their rank *r* as a Rank Score (RS), computed as *n-r+1* in each network, where a higher rank score indicates higher centrality. RS are weighted (WRS) by an exponent *p* to give more significance to differences between high rankings rather than low rankings where *WRS* = *rS^p^* and *p* = 25 in the Pan-Cancer analysis. WRS are normalized (NWRS) for each network where *NWRS = wrs/sum(wrs*) within a single network. Using the NWRS, a different centrality score (DCS) for each node is computed as the *max(wnrs)-mean(nwrs*) across networks. We sorted genes by their DCS to find genes differentially central in one or few networks relative to the rest.

### Enrichment of Tumor Survival Genes

We selected the top and bottom 100 genes ranked by their eigen-centrality in each network and visualized their dependency score across various cell lines from the Cancer Dependency Map Project (DepMap) through the R package *depmap* (v1.2); specifically using the CRISPR - Achilles gene effect resource (EH2261) as well as associated cell line metadata (EH2266)^37^. Genes ranked within the top and bottom 100 for each network that were not present in the DepMap were omitted from the analysis. We performed a Wilcoxon test in each network to assess the significance in difference between the distributions of dependency scores for genes within the top and bottom centrality groups.

### Network Propagation

We used the random walk with restart (RWR) algorithm, which measures the distance (or proximity) of nodes in a graph from one or multiple seed nodes^41^. It does this by randomly traversing the graph with a given restart probability, starting from the seed node(s). We used the proximity values to visualize the graphical distance of nodes of interest to other nodes in the graph. To characterize the biological function of a set of seed nodes, we ranked traversed nodes by their proximity to the seed nodes(s), which is used as input to the ranked-based Kolmogorov-Smirnov test for enrichment implemented in the R package *hypelR*^55^.

### Essential Genes Signaling Pathways

We performed the described network propagation technique (*restart* = 0.5) using the 14 eigen-central genes as seed nodes for each breast cancer subtype network. In Fig. 5C we use the parent network (L2) to visualize the network propagation which had a similar global structure to subtype networks (L3). Enrichment was performed using the Reactome curated genesets from MSigDB v7.2.1 and filtering by FDR < 0.001. Because many of the genesets contain similar gene members and are redundant, we used a hierarchical clustering method based on a pairwise Jaccard index to selectively highlight non-redundant genesets enriched across multiple networks^56^.

### Biological Pathway Enrichment

Enrichment of biological processes and pathways was performed with the R package *hypeR*. We used both the hypergeometric test for overrepresentation and the ranked-based Kolmogorov-Smirnov test for enrichment. We used curated genesets from MSigDB including v7.2.1 of the Hallmark and Reactome collections.

### Disease Neighborhoods

We performed community detection on networks using the Walktrap^38^ algorithm (*steps* = 10). Communities were tested for enrichment of known breast cancer disease genes via a hypergeometric test as previously described and using expert curated breast carcinoma disease genes (C0678222) from DisGeNET^39^ (v7.0). Identified disease neighborhoods were tested for enrichment of biological pathways using the hypergeometric test for overrepresentation with the R package *hypeR*. Enrichment was performed using the Hallmark curated genesets from MSigDB v7.2.1 and filtering by FDR < 0.05. Guilt by association with breast cancer disease genes was computed by performing network propagation within each disease neighborhood, using disease genes as seed nodes as previously described (*restart* = 0.7). Druggable targets were identified from The Genomics of Drug Sensitivity in Cancer^43^.

## DATA AVAILABILITY STATEMENT

The presented methods are available as an open source R package named *shine*. The package supports high-dimensional constraint-based structure learning for network hierarchies. We have additionally made available the Nextflow-based framework for inferring multiple large-scale networks on high performance computing environments called *shine-nf*. The learned networks have been published to the Network Data Exchange (NDEx) and are also hosted through our publicly accessible interactive web portal.

Portal: bieulergy.shinyapps.io/shine

Repositories: github.com/montilab/{shine,shine-nf}

Documentation: montilab.github.io/shine

Operating system: Linux, OS X

Programming languages: R, Nextflow

License: GNU GPLv3

## ACKNOWLEDGMENTS

This work was supported by the Find the Cause Breast Cancer Foundation, the National Cancer Institute (NCI U01CA243004), the National Institute on Aging (NIA cooperative agreement UH2AG064704), the Moorman-Simon Fellowship in Computational Biomedicine, as well as the National Institute of Dental & Craniofacial Research of the National Institutes of Health under Award Number F31DE029701. The authors would like to thank R. Mohammadi, author of the R package BDgraph, for his assistance with inquiries related to Bayesian structure learning through his software.

## DECLARATION OF INTERESTS

The authors declare no competing interests.

## Notes

### Competing Interest Statement

The authors have declared no competing interest.

